# Impacts of early exposure to ethanol on adenosine functioning in zebrafish

**DOI:** 10.1101/2023.08.03.551856

**Authors:** Giovanna Trevisan Couto, Guilherme Pietro da Silva, Liliana Rockenbach, Jéssica Scheid da Silva, Monica Ryff Moreira Roca Vianna, Rosane Souza Da Silva

**Author notes:** Corresponding author: Rosane Souza Da Silva, Programa de Pós-Graduação em Neurociências Universidade Federal, Fluminense – Campus do Gragoatá, R. Alexandre Moura, 8 – Bloco M – Instituto de Biologia São Domingos, Niterói, – Rio de Janeiro – CEP: 24210-200.

## Abstract

Exposure to ethanol at the beginning of development can impact the formation of the Nervous System. The set of symptoms resulting from ethanol consumption during pregnancy is called FASD (Fetal Alcohol Spectrum Disorders) and ranges from cognitive alterations to the most severe form called FAS (Fetal Alcohol Syndrome). The effect caused by ethanol on the formation of brain architecture directly affects the adenosine neuromodulation system. In this work, a single exposure regimen of 24 to 26 hpf to 1% ethanol transdermally was used as a model to assess adenosine signaling in the context of seizure susceptibility in zebrafish larvae and adults. To test sensitivity, a 2.5 mM subconvulsant dose of pentylenetetrazole (PTZ) was used, which was not able to increase seizure events in larvae or adults exposed to ethanol during embryonic phase. However, the duration of stage I was increased and the latency to reach stage II was decreased in larvae, showing a possible proconvulsant profile in these ethanol-treated animals. Also, the exposure of larvae to CPA (75 μM) was able to reverse the effect of embryonic ethanol treatment on the latency to reach stage II of seizure. Adenosine and ecto-5’-nucleotidase receptor mRNA expression did not show significant difference in both developmental stages. These results demonstrated that even a short and specific exposure to ethanol can promote, even if mild, effects on neuronal modulation, increasing susceptibility to seizures.

## 1. Introduction

Fetal Alcohol Spectrum Disorder (FASD) is an important challenge for public health management, especially in developing or underdeveloped countries (1, 2). It is estimated that more than 1,700 new cases of FASD are identified globally each day (2). FASD includes severe forms, such as Fetal Alcohol Syndrome (FAS), which is characterized by morphological changes in the face, reduced body size, auditory and visual problems, among others, which is easily diagnosed (3). Milder forms, called alcohol-related neurodevelopmental disorders (ARND), exhibit cognitive changes even when morphological sequelae are absent (4). Hyperactivity, attention difficulties and damage to learning and memory are some of these alterations (5).

Ethanol exposure is a risk factor for the development of structural and functional abnormalities, impacting neuronal survival, migration, cell differentiation and synaptogenesis (6). Additionally, studies indicate that 3 to 21% of individuals with FASD develop some type of convulsive state ranging from electroencephalographic changes to epilepsy (7). The development of such sequelae may be dependent on the gestational period and exposure dose. Human studies suggest that short-term, large-volume ingestions of ethanol during gestational weeks 11–16 increase the risk of neonatal seizures and the risk of developing epilepsy (8).

Subsidies for understanding these impacts include the ability of ethanol to interact with various proteins, such as glutamatergic, serotonergic, glycinergic, GABAergic and purinergic receptors, in addition to ion channels (9). Among these targets, the purinergic system stands out, given the importance of adenosine for neuromodulation, for the brain development, and for seizure response (10). In animal studies, acute ethanol exposure raises extracellular levels of adenosine by inhibition of equilibrative nucleoside transporters (ENT1), indirectly stimulating adenosine P1 receptors (11, 12). Previous data from our group indicated that in a FASD model in zebrafish there is an elevation of brain ecto-5’-nucleotidase enzyme activity which is persistent into adulthood (13). Furthermore, this model of FASD in zebrafish can promote cognitive alterations that mimic human sequelae, such as memory deficits and impairment of sociability, which were prevented by the administration of ecto-5’-nucleotidase inhibitors (14, 15). Therefore, in this work, we sought to investigate the participation of adenosinergic modulation in the possible long-term effects of ethanol given at the beginning of development on the sensitivity to seizures through the pentylenetetrazol-mediated seizure sensitivity measure and the analysis of the response to the anticonvulsant effect of an adenosine A_1_ receptor agonist. Also, we performed an analysis of gene expression of high affinity adenosine P_1_ receptors and the enzyme ecto-5’-nucleotidase at the timepoint of seizure sensitivity evaluation.

## 2. Methodology

### 2.1 Drugs

Ethanol was obtained from Neon (Sao Paulo). CPA (Sigma Aldrich - ref: C8031) Pentylenetetrazole (Sigma Aldrich - ref P6500) and Tricaine (MSS-222) were purchased from Sigma Chemical Co. (St. Louis, MO, USA). Trizol was Life Technologies USA (Reference - 15596026).

### 2.2 Animal maintenance

Zebrafish (*Danio rerio*) of the wild lineage AB was bred and kept in the institutional vivarium Center for Biological and Experimental Models (CEMBE – PUCRS) in an automated recirculating system (Zebtec, Tecniplast, Italy). This system used water filtered by reverse osmosis at a temperature of 26.5± 1.5°C, pH 7-7.5, oxygenation at 7.20 mg O_2_/L, and conductivity at 300–700 μS. The light / dark cycle was 14h/10h. The animals were fed from 6 days post-fertilization (dpf) with paramecium and ground feed (TetraMin Tropical Flakes Fish) and from 13 dpf artemia was added (16). Adults were fed three times a day with flakes and brine shrimp. Fifteen adult mating matrices animals at a proportion of 2 males to 1 female produced the embryos used in these experiments. The fertilized eggs were collected at 30 minutes and randomized assigned for control or ethanol group. After treatment animals followed the standard maintenance procedure described above. The animals used for seizure evaluation were euthanized by anesthetic overdose (buffered tricaine solution, 0.5 g.L^-1^) at the end of the experiments. Those animals that underwent tissue extraction for gene expression analysis were euthanized by hypothermic shock in flocculated ice water. All manipulation of animals were in accordance with the procedures described by Westerfield (16) and followed the guidelines of the Brazilian National Council for the Control of Animal Experimentation (17). All protocols were approved by the Institutional Animal Care Committee of the Pontifical Catholic University of Rio Grande do Sul (CEUA/PUCRS 9570/2019).

### 2.3 Exposure to ethanol

Embryos (24 hours post-fertilization (hpf)) were transdermally treated for 2h with a 1% (v/v) ethanol solution diluted in water from the system and kept in a Petri dish (8.7 cm diameter). After this period, the embryos were rinsed and kept into water free of drug from the system. All embryos were kept at 28 ± 1°C in a Bio-Oxygen Demand incubator with light/dark cycle of 14/10 h until they are used for the experiments at 7 dpf or transferred to growth aquariums in automated racks with recirculating system water until they reach 3 months post-fertilization (mpf) in the standard conditions described in item 2.2. Control animals underwent the same manipulations, except for exposure to ethanol, being exposed exclusively to water from the system.

### 2.4 Assessment of seizure susceptibility

#### 2.4.1 Larval zebrafish

At 7 dpf, zebrafish larvae from both experimental groups were subjected to a subconvulsant-dose of Pentylenetetrazole (PTZ) (2.5 mM; Couto et al., 2023) to verify seizure sensitivity. PTZ exposure was performed individually in a 24-well cell culture plate (1.6 cm well diameter) containing 3 mL of PTZ solution each well. Filming was performed during PTZ exposure (10 minutes) by a digital camera (Logitech, Romanel-sur-Morges, Switzerland) located at the top of the Ethovision XT apparatus (Noldus, Netherlands). All experiments were carried out between 10 am and 3 pm. Each experimental group of larvae was of 16 animals, which was calculated considering a power effect of 80 %. Data were extracted using Ethovision XT 8 Software Package (Noldus, Netherlands). Seizure stages followed those described by Baraban (18). Briefly, stage I marked by a drastic increase in swimming activity, stage II by circle swimming, and stage III by clonus-like seizures with loss of posture and immobility. The parameters analyzed were frequency of animals reaching each stage, the latency (seconds) to reach each stage and the duration (seconds) of seizure in each stage. The analyzes of the recordings were made blindly by two observers.

#### 2.4.2 Adult zebrafish

At 3 mpf, another set of animals were acclimated in the experiment room 24 hours before the analyzes and without food. Individual exposure to PTZ (2.5 mM) occurred in aquariums (13 x 11.5 x 8 cm; height x length x width) containing 500 mL of the PTZ solution (water column of 5.5 cm in height). Immediately, filming began and lasted for 6 minutes using a frontal digital camera (Logitech, Romanel-sur-Morges, Switzerland) at 65 cm. All experiments were carried out between 10 am and 3 pm. The sample number was 12 animals (19). Data were extracted using Ethovision XT 8 Software Package (Noldus, Netherlands). The seizure stages considered for the adult animals were stage II characterized by an increase in speed, erratic movements to the right and left, stage III by increased speed and circular movements, stage IV by abrupt increase in speed, muscle contractions with jumps and abrupt movements, and stage V by loss of posture, falling to the bottom of the aquarium and body rigidity (20, 21). The number of animals reaching each seizure stage and the duration (seconds) and latency (seconds) to reach each seizure stage were recorded. The analyzes of the recordings were made blindly by two observers.

### 2.5 Anticonvulsant action of adenosine A_**1**_ receptor agonist (CPA)

Concomitantly with the PTZ sensitivity analysis, an additional experimental group was used to evaluate the anticonvulsant action of CPA (N_6_-Cyclopentyladenosine). Prior to exposure to PTZ, this experimental group, which was maintained under the conditions described in items 2.2 and 2.3, was exposed to CPA. The 7 dpf larvae were exposed for 30 minutes to a 75 μM CPA solution in a 24-well plate with 3 ml each (22). For adult animals intraperitoneal injection of CPA (10 mg.Kg^-1^) was done 30 minutes before exposure to PTZ using a microsyringe (NanoFil – World Precision) in a volume of 5 μL (10 mL.Kg^-1^)(22). The animals were previously anesthetized in groups of 4 in a beaker with 300 ml of solution containing 0.03 g of tricaine and immediately after the identification of decreased swimming, stagnation, loss of muscle tone and belly turning upwards, received CPA injection or saline (Controls). The number of animals per group and all protocols were the same described at items 2.2 to 2.4.

### 2.6 Analysis of relative gene expression of P1 receptors and ecto-5’-nucleotidase enzyme by qPCR

To evaluate the effect of exposure to 1% ethanol on the expression of the *adoraab, adoraa2aa, adoraa2ab* and *nt5e* genes, 7 dpf and 3 mpf animals, free from PTZ interference, were euthanized in an ice bath (23). The sample size was 4 experimental unit per experimental group (control and ethanol). Each experimental unit represents a pool of 20 larvae, totaling 80 larvae per experimental group or a pool of 3 brains totaling 12 adult animals per experimental group. In plastic microtubes (1.5 mL), each sample received 300 μl of trizol and these were stored in a freezer at -80°C until the day of the analysis (up to a week). For mRNA extraction, frozen samples were previously macerated and homogenized with vortex. Chloroform was added for phase separation. After precipitation with isopropanol, they were homogenized and centrifuged, with the supernatant discarded, conserving only the pellet. For washing, 75% ethanol was used, and the sample homogenized and centrifuged, and only the pellet was conserved again. Finally, the samples were resuspended in RNAse-free water and RNAse OUT was added to preserve the final volume. RNA quantification was performed using a NanoDrop spectrophotometer (Thermo Scientific - NanoDrop Lite Printer) from the determination of the absorbance ratio at 260 nm and 280 nm using an aliquot of 1 μL of each sample for reading. After digestion with DNAse I (Sigma Kit 1002268809) to purify the RNA, the High – Capacity cDNA Reverse Transcription kit (Thermo Fisher) was used to cDNA synthesis with 1 μg of RNA, following all manufacturer’s instructions. The relative expression of the genes listed in Table 1 was determined by quantitative reverse transcription polymerase chain reaction (qRT-PCR) using PowerUp SYBR® Green (Invitrogen) and the Line Gene 9660 Thermal Cycler (BIOER). Beta actin and EF1-alpha were used as reference genes for normalization and determination of relative mRNA expression levels.

**Tabela 1:**
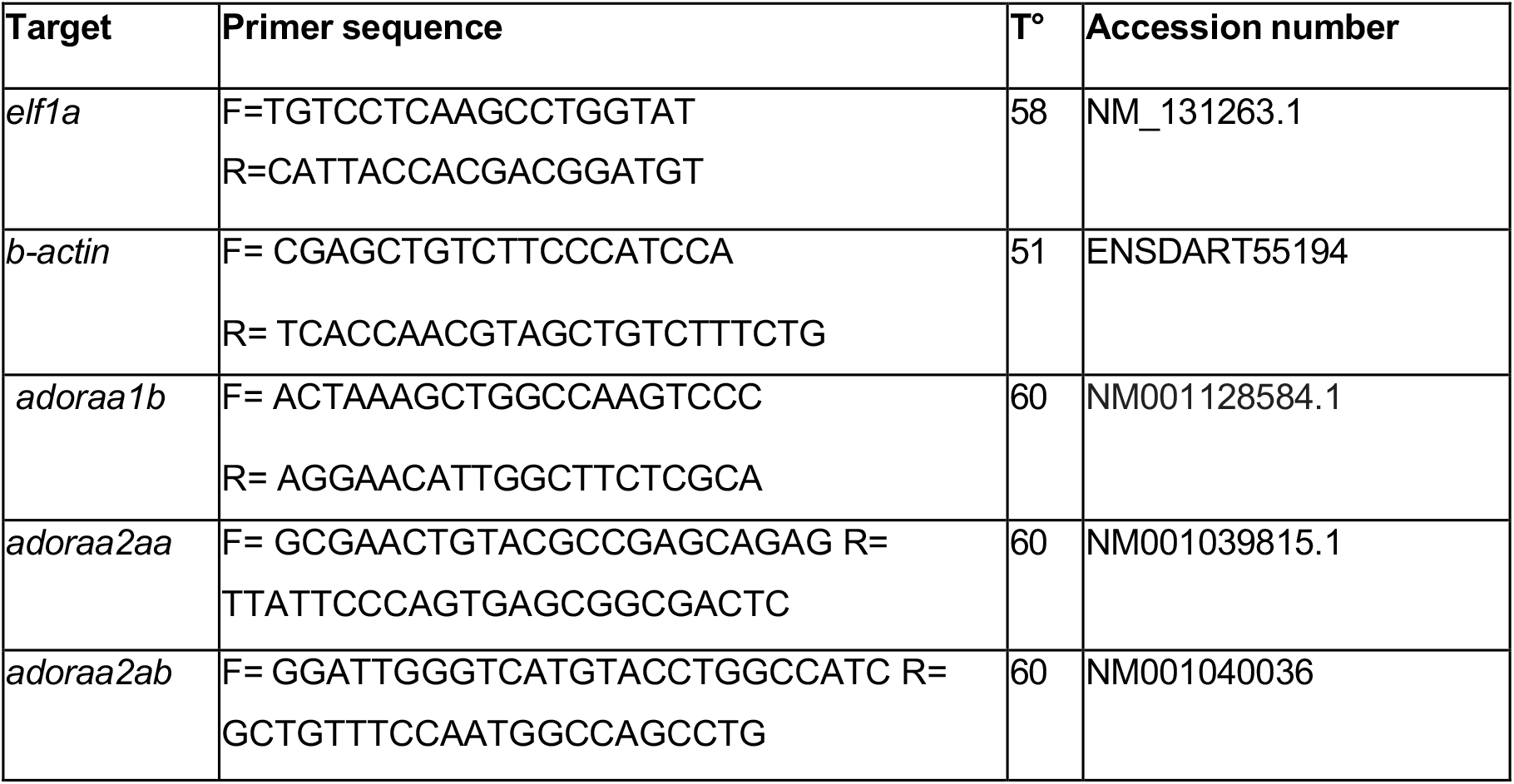
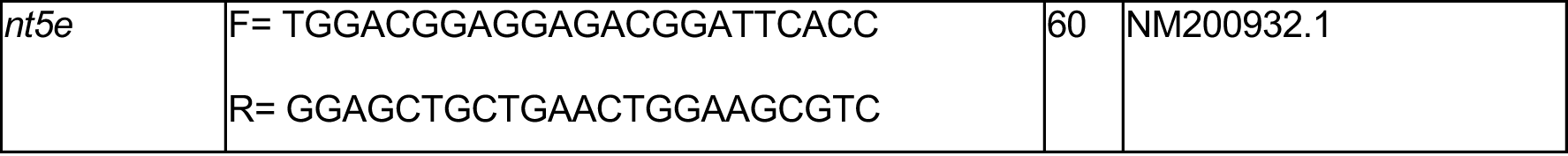
Sequência de primers dos genes alvo (14, 24).

### 2.7 Statistical analysis

For seizure assessment, data were presented as frequency of animals reaching specific seizure stages. Comparison between frequencies was performed using the Chi-square test. For those groups with more than 3 animals reaching the seizure stages, the mean latency and duration of the convulsive stages were analyzed using the Kruskal-Wallis test and the data presented as mean ± standard error of the mean. Relative gene expression was compared between control and treated groups using the Mann-Whitney test. Statistical differences were attributed when p<0.05.

## 3. Results

### 3.1 Sensitivity to Pentylenetetrazol

The frequency of larvae reaching convulsive stages I and II did not differ between the animals in the control group and the group treated with ethanol, in the same way that pre-treatment with CPA did not change this outcome (p=0.4096, Chi test - square) (Figure 1A). However, the ethanol group reached significantly lower latencies to reach stage I than those that received pre-treatment with CPA (Fig. 1B) (p=0.046, Kruskal-Wallis Test), although these groups did not differ from control animals (Figure 1B). The duration of seizure stage I for larvae subjected to ethanol in the embryonic phase was significantly longer than for control animals, while previous treatment with CPA was able to alter this significant effect in comparison to ethanol group but remained different from control animals (p=0.006; Kruskal-Wallis Test).

**Fig. 1.**
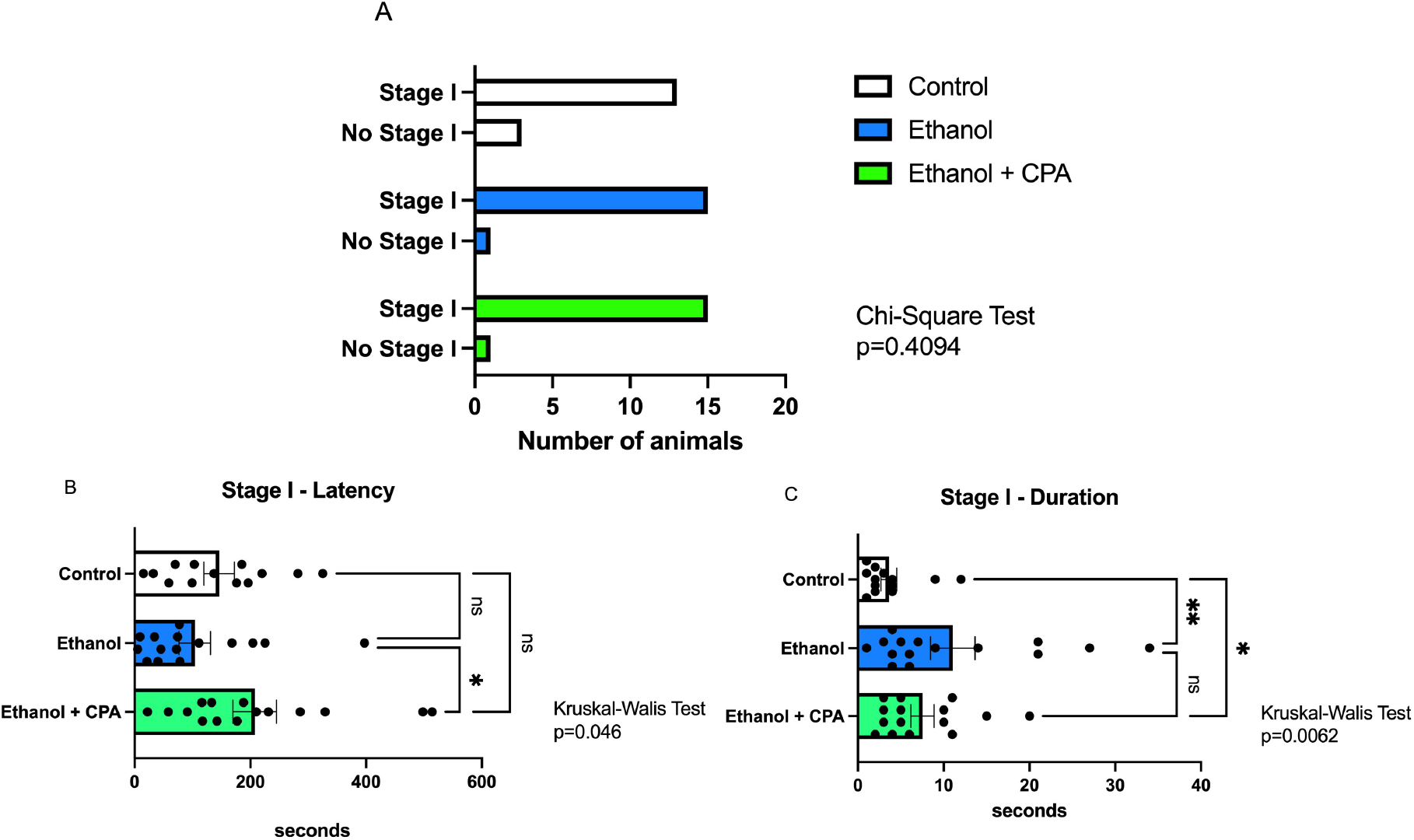
Evaluation of susceptibility to stage I seizures induced by PTZ in 7 dpf larvae exposed to ethanol (1% v/v) in the embryonic stage (24-26 hpf). A) Frequency of larvae that reached stage I on the seizure scale for the control, 1% ethanol treated and 1% ethanol + CPA treated groups. B) Latency in seconds to reach seizure state I; C) Duration in seconds in stage I. CPA was given transdermally at a concentration of 75 μM, 30 minutes before exposure to PTZ (2.5 mM; 10 minutes). Controls were only exposed to water. Sample number=16 larvae per experimental group.

The frequency of animals undergoing stage II was not affected by treatments (p=0.7058, Chi-square test) (Figure 2A). The latency to reach stage II was significantly lower in animals exposed to ethanol in the embryonic phase than in controls while CPA treatment was able to change this outcome but not reaching control levels (Figure 2B) (p=0.03, Kruskal-Wallis Test). The duration of stage II did not differ between the experimental groups (Figure 2C) (p=0.6845, Kruskal-Wallis test). There were no larvae reaching seizure stage III.

**Fig. 2.**
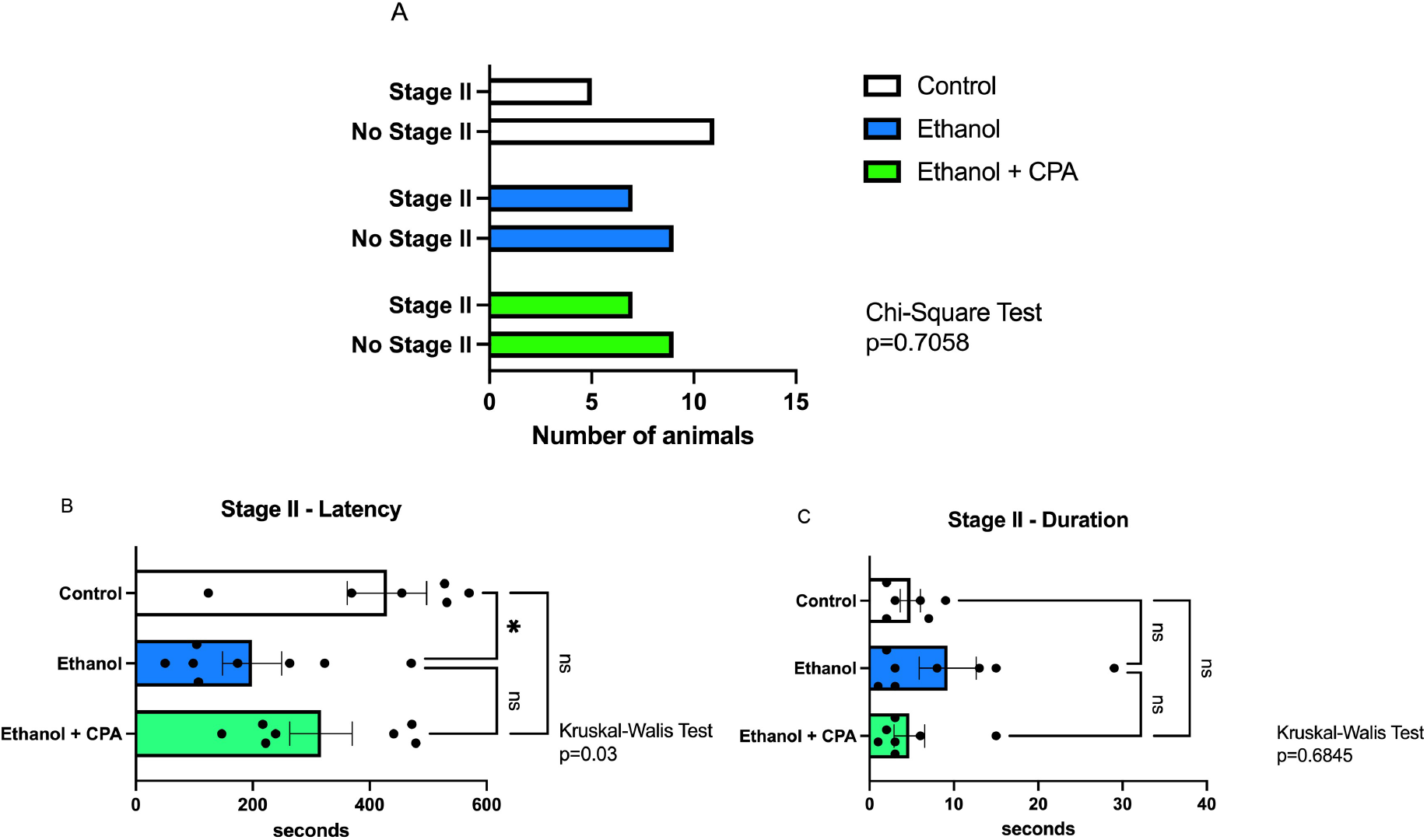
Evaluation of susceptibility to stage II PTZ-mediated seizure in 7 dpf larvae exposed to ethanol (1% v/v) in the embryonic stage (24-26 hpf). A) Frequency of larvae that reached stage II in the seizure scale for the control, 1% ethanol treated and 1% ethanol + CPA treated groups. B) Latency in seconds to reach seizure state I; C) Duration in seconds in stage I. CPA was given transdermally at a concentration of 75 μM, 30 minutes before exposure to PTZ (2.5 mM; 10 minutes). Controls were only exposed to water. Sample size =16 larvae per experimental group.

Most adult animals reached seizure stage II, while the minority reached stages III and IV, but the frequency of adult animals reaching these stages did not differ between the experimental groups (II: p=0.1019; III: p=0.1286; IV: p=0.225, Chisquare Test) (Figure 3A, B, C). For animals that reached stage II, both the latency and the duration of this stage did not differ between the experimental groups (p=0.1954; p=0.6158, respectively, Kruskal-Wallis Test) (Figure 3D and E).

**Fig. 3.**
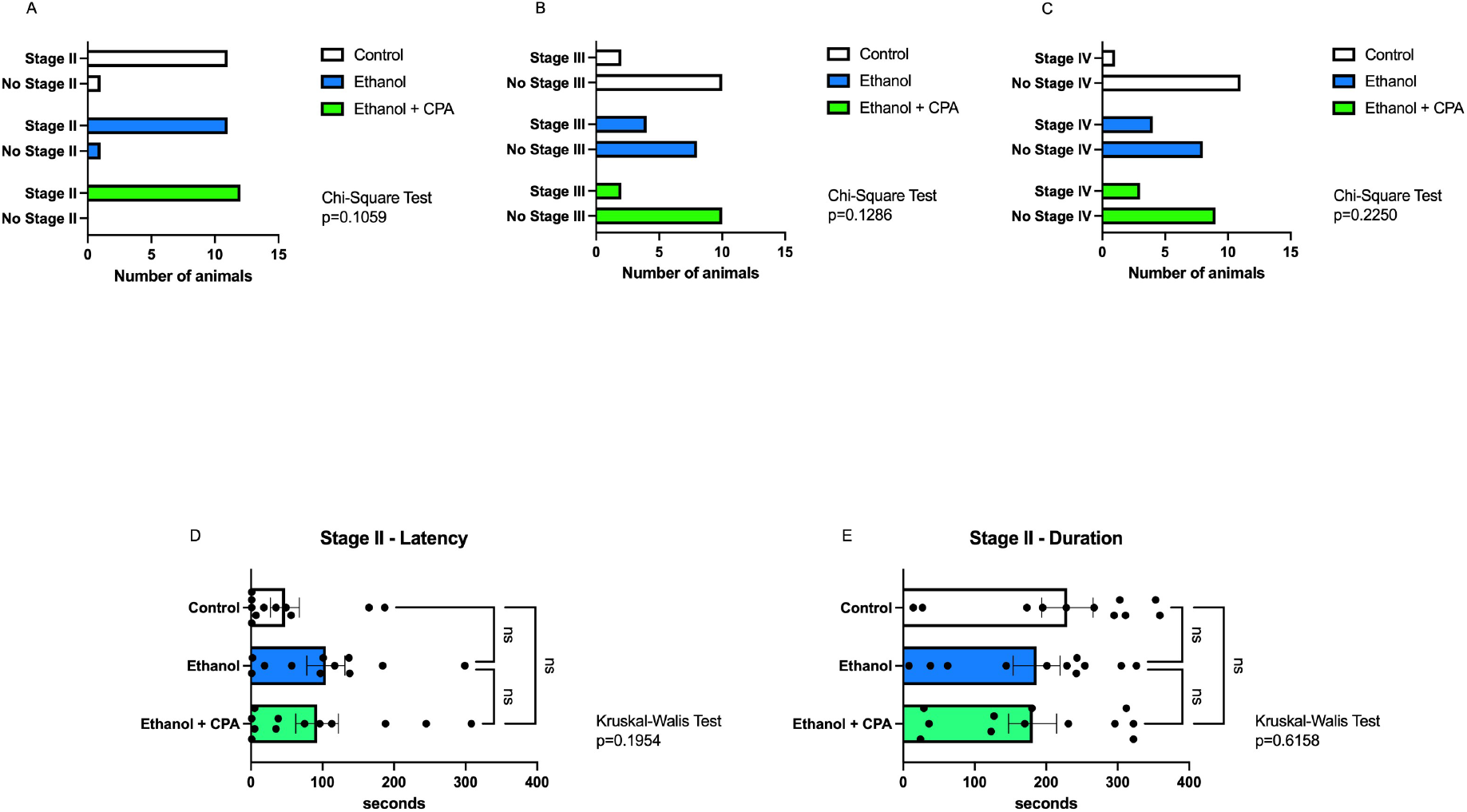
Evaluation of susceptibility to PTZ-mediated seizure stages II, III and IV in adult zebrafish (3 mpf) exposed to ethanol (1% v/v) in the embryonic stage (24-26 hpf). A, B and C) Frequency of animals that reached stages II, III and IV on the seizure scale, respectively, for the control, 1% ethanol and 1% ethanol + CPA groups. D) Latency in seconds to reach state II of seizure; E) Duration in seconds in stage II. CPA was given intraperitoneally at a dose of 10 mg. Kg-1, 30 minutes before exposure to PTZ (2.5 mM; 6 minutes). Controls were only exposed to water. Sample size =12 animals *per* experimental group.

### 3.2 Analysis of relative gene expression of adenosine receptors and ecto-5’-nucleotidase by qPCR

The expression of the genes that code for the A1, A2AA and A2AB receptors (*adoraa1b, adoraa2aa*, and *adoraa2ab*) was analyzed together with the main enzyme producing adenosine in the extracellular medium, the ecto-5’-nucleotidase (*nt5e*) in animals of 7 dpf and 3mpf exposed to ethanol in the embryonic phase (24-26 hpf). Both the evaluation of expression in whole larvae and in adult zebrafish brain did not differ in the experimental groups for any of the evaluated genes (Larvas: *adoraa1b*: p= 0.7429, *adoraa2aa*: p=0.9714, *adoraa2ab*: p=0.6857; *nt5*e: p=0.8857; Adults: *adoraa1b*: p= 0.4857, *adoraa2aa*: p=0.4, *adoraa2ab*: p=0.8571; *nt5e*: p=0.6286, Mann-Whitney test) (Figures 4 and 5).

**Fig. 4.**
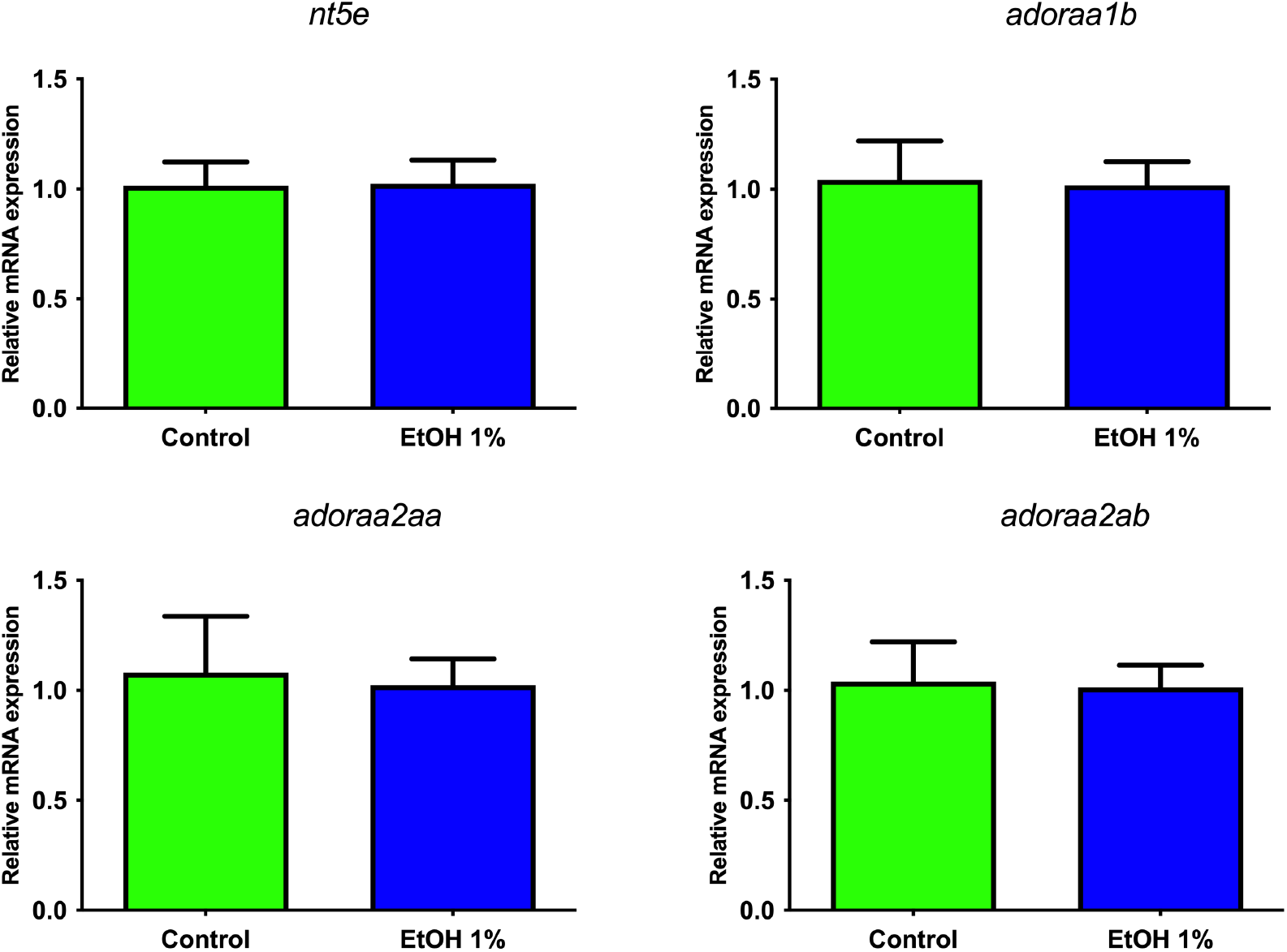
Relative gene expression of adenosine receptor and ecto-5’-nucleotidase mRNA in whole larvae (7dpf) after embryonic exposure (24-26 hpf) to 1% ethanol (v/v). Controls were only exposed to water. *nt5e*= ecto-5’-nucleotidase; *adoraa1b*-A1 receptor; *adoraa2aa*: A2A1 receptor; and *adoraa2ab*: A2A2 receptor. Sample size = 4 experimental unit per experimental group. Mann-Whitney statistical analysis.

**Fig. 5.**
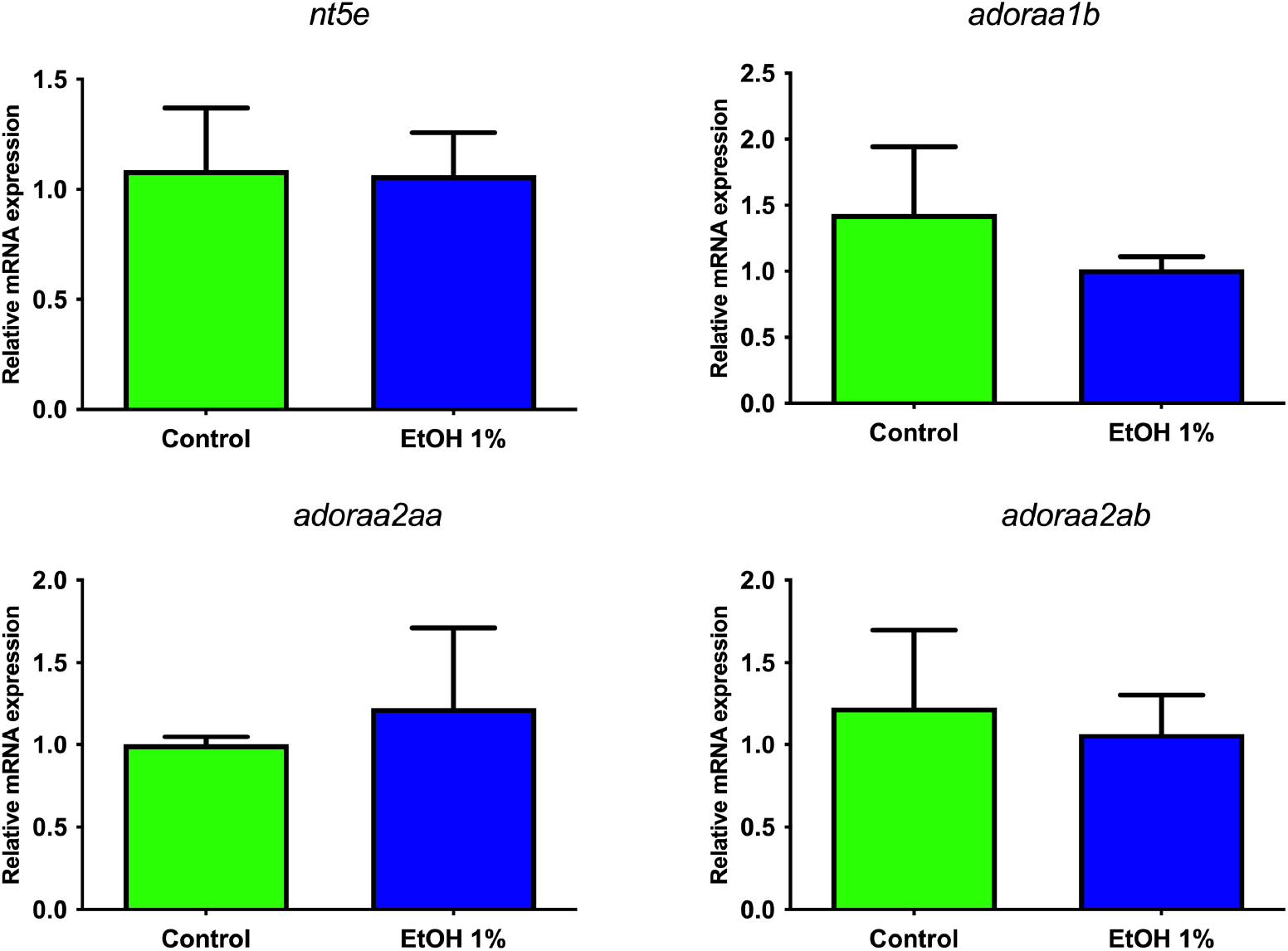
Relative gene expression of adenosine receptor and ecto-5’-nucleotidase mRNA in adult zebrafish brain (3mpf) after embryonic exposure (24-26 hpf) to 1% ethanol (v/v). Controls were only exposed to water. *nt5e*= ecto-5’-nucleotidase; *adoraa1b*-A1 receptor; *adoraa2aa*: A2A1 receptor; and *adoraa2ab*: A2A2 receptor. Data are presented as mean ± SEM. Sample number =. 4 *per* experimental group. Mann-Whitney statistical analysis.

## Discussion

Although studies indicate that acute exposure to ethanol can alter the availability of adenosine in the extracellular environment and the functionality of its specific receptors (11, 12, 25), it is still unknown whether these alterations, when early in development, can lastingly affect CNS functioning, in particular contributing to susceptibility to seizures, given the natural anticonvulsant role of adenosine .

Here, we exposed zebrafish embryos to ethanol in the period from 24 to 26 hpf, which means in the pharyngeal phase (24-48 hpf). At this stage of zebrafish development, the body begins to organize dorsally, becomes vascularized and begins to organize itself to form the Nervous System (26). In vertebrate general development, this stage is an intense period of migration and cell differentiation until the beginning of brain development (27, 28). The 1% concentration used in this study is prevalent among studies with exposure of zebrafish to ethanol, as it reproduces neurochemical impacts without affecting the animal’s morphology (29). Although in direct comparison with the exposure of human embryos to ethanol this 1% seems high, the barrier exerted by the chorion in zebrafish embryos must be considered, since the concentration of ethanol inside the egg is 2.7 to 6.2 times lower than the external concentration used (30).

Here, we used a subconvulsant concentration of PTZ, which worked for the strongest and most representative seizure stages (II for Larvae and IV for adults), while for stages more based on the locomotion status (I for Larvae and II for adults) this PTZ concentration was able to increase the locomotion of animals. Despite that, no difference in the frequency of animals reaching all investigated convulsive stages, including the most characteristic ones, both for larvae (stage II) and adults (stage IV), when compared to control group. Although, this result may indicate that the exposure to ethanol did not have any effects on seizure sensitivity, it is also possible to consider that, at least for the larval stage, the PTZ concentration was insufficient. Zebrafish larvae have been found to be more resistant to PTZ, and concentrations such as 15 mM are often used to achieve seizure (18). In the other hand, in a similar approach by Wang et al. (31), zebrafish larvae treated with ethanol during embryonic period had decreased latency to reach convulsive stages in a dose dependent manner of PTZ. In that report, even the 2.5 mM concentration of PTZ was able to induce convulsive response at stage I and II that was affected by ethanol pretreatment, while stage III was not reached (31). These differences could be a matter of genetical background of animals since the susceptibility to PTZ seems different but is worth to mention that the stage III was not reached during exposure to 2.5 mM in both studies. In relation to ethanol effects, the comparison between these two studies emphasizes the importance of timing in ethanol exposure, since our study was more restrictive (24-26hpf) and Wang study used an expanded period of exposure (3-24hpf).

For larvae reaching I and II seizure stages, it was possible to explore the latency and duration to reach such stages. The duration of stage I was longer for larvae that received ethanol in the embryonic phase in relation to controls, while the latency to reach this stage appears reduced, but not reaching statistical significance. The latency for stage II was lower, while the duration also seems longer but with no statistical significance. This may suggest a greater sensitivity to seizures on the ethanol-treated animals, as we used subconvulsant doses of PTZ. The frequency of adult animals reaching seizure stages II, III and IV was not different between control and ethanol-treated animals. The only stage with enough animals for the analysis of latency and duration of stage was the stage II. No statistical significance was attributed to these parameters.

Although the data for PTZ sensitivity did not differ sharply between ethanol-treated and control animals, the use of the rigid adenosine agonist, CPA, showed effects in increasing the latency to stage I in larvae compared to ethanol-treated group. Literature findings proved that exposure to ethanol in the embryonic stage of zebrafish increases the activity of ecto-5’-nucleotidase (25), promotes a robust reduction in serotonin and dopamine levels (32), increased number of apoptotic cells and persistent decrease in glutamate transporters (33, 34).These alterations have been related to cognitive deficits that resemble those found in human individuals who have FASD (14, 15, 29, 35-37).

This possible differential response of ethanol-treated animals to CPA could be related to long-lasting effects on adenosine receptor expression, binding, or trafficking. Here we can see that the gene expression of A_1_ and A_2_ receptors and ecto-5’-nucleotidase from total larvae and brain from adult animals was not affected by embryonic exposure to ethanol.

Under the conditions tested here, embryonic exposure to ethanol contributed slightly to seizure susceptibility, and receptors and ecto-5’-nucleotidase did not have their gene expression affected in the short and long term. This may be in line with the fact that the present study mimicked a mild within-spectrum form of fetal alcohol syndrome with a short, single exposure to ethanol. It is noteworthy that even this short and specific exposure to ethanol can promote, even if mild, effects on convulsive parameters that do not last until adulthood. It remains to be investigated whether, in models closer to more severe forms of FASD, these and other biological components underlying adenosine availability and functionality through P1 receptors, such as enzymatic activity of adenosine deaminase, nucleoside transport mechanisms, and adenosine ligand-receptor binding capacity, contribute to increased susceptibility to seizures.

## Consent for Publication

All authors agreed to this publication.

## Acknowledgements

We thanks to Fernanda Morrone for the support in PCR analysis.

